# Persistent vulnerability to heroin relapse across the adult lifespan in rats

**DOI:** 10.64898/2026.03.18.712140

**Authors:** Rajtarun Madangopal, Olivia R. Drake, Diana Q. Pham, Veronica A. Lennon, Sophia J. Weber, Justine Lee, Adekemi Sobukunola, Ava R. Holmes, Omodolapo Nurudeen, Yavin Shaham, Bruce T. Hope

## Abstract

Relapse to opioid use during abstinence is often triggered by drug-associated cues but the persistence of this effect across the lifespan is unknown. Using a rat model, we found that relapse provoked by heroin-predictive discriminative stimuli persisted for over one year of abstinence, suggesting enduring, potentially lifelong opioid relapse vulnerability.

## Introduction

Relapse is a core feature of opioid addiction, and relapse after prolonged abstinence is often triggered by craving-inducing cues and contexts previously associated with opioid use [1]. Human studies indicate that cue-induced craving can persist for up to 2-3 years [2,3], but whether vulnerability to craving and relapse extends across the adult lifespan is unknown. Addressing this question in humans is neither practical nor ethical as it would require random assignment to different abstinence durations prior to cue exposure [4].

Animal models allow investigation of persistent relapse vulnerability by measuring responses to opioid-paired cues or contexts after extinction of previously drug-reinforced responding at varying abstinence intervals [5], or following extended experimenter-imposed or voluntary abstinence [3]. Using these approaches, prior studies have demonstrated persistent cue- and context-induced opioid-seeking for up to 2 months of abstinence [6]. However, as in humans, whether relapse vulnerability persists across the adult rat lifespan is unknown.

Based on prior cocaine studies showing enduring discriminative stimulus (DS) control over drug-seeking [7,8], we developed a trial-based discrimination procedure that isolates DS contributions from contextual and drug-paired conditioned stimuli (CSs) [9]. We showed that DS-controlled relapse to cocaine-seeking progressively increases (“incubates”) up to 60 abstinence days and persists up to 300 days. Here, we used this procedure to assess lifespan relapse vulnerability following repeated exposure to DSs signaling heroin availability (DS+) and unavailability (DS−). We also tested whether DSs retain control over heroin-primed relapse after over one year of abstinence.

## Methods

We trained rats to self-administer heroin under discriminative stimuli signaling heroin availability (DS+) or unavailability (DS−). We then repeatedly tested DS-controlled relapse to heroin-seeking at different abstinence days for over 1 year. To isolate the contribution of each DS, we independently varied DS+ and DS− presentation (on/off) in a four-condition experimental design (no DSs, DS+ only, DS− only, both DSs) and assessed heroin-seeking after 3 weeks of abstinence. We also evaluated DS control of heroin-primed reinstatement after prolonged abstinence. Detailed methods and analyses are provided in Supplementary Online Methods.

### DS-controlled heroin-seeking during prolonged abstinence

In Experiment 1 (Figure 1A), we examined the timecourse of DS-controlled heroin-seeking across over 1 year of abstinence and tested whether heroin-priming reinstates DS-controlled heroin-seeking. Following heroin self-administration training, we trained rats (n=13 male, 13 female) to press a central retractable lever during trials where a light-cue preceded lever insertion and signaled heroin availability (DS+ trials), and to withhold responses during trials where a different light-cue preceded lever insertion and signaled heroin unavailability (DS− trials). Each 3-h discrimination training session included 30 DS+ and 30 DS− trials, and lever-presses during DS+ trials were reinforced with heroin (0.025 mg/kg/infusion; no reward-paired discrete cues were available). After training, we assessed relapse to heroin-seeking in 3-h extinction tests using the same DS trial structure after 1, 21, 60, 120, 200, 300, and 385 abstinence days (within-subjects). After the relapse tests, we conducted one additional 3-h extinction session to extinguish DS+ responding and tested whether heroin-priming injections reinstate DS-controlled heroin-seeking.

**Figure 1:**
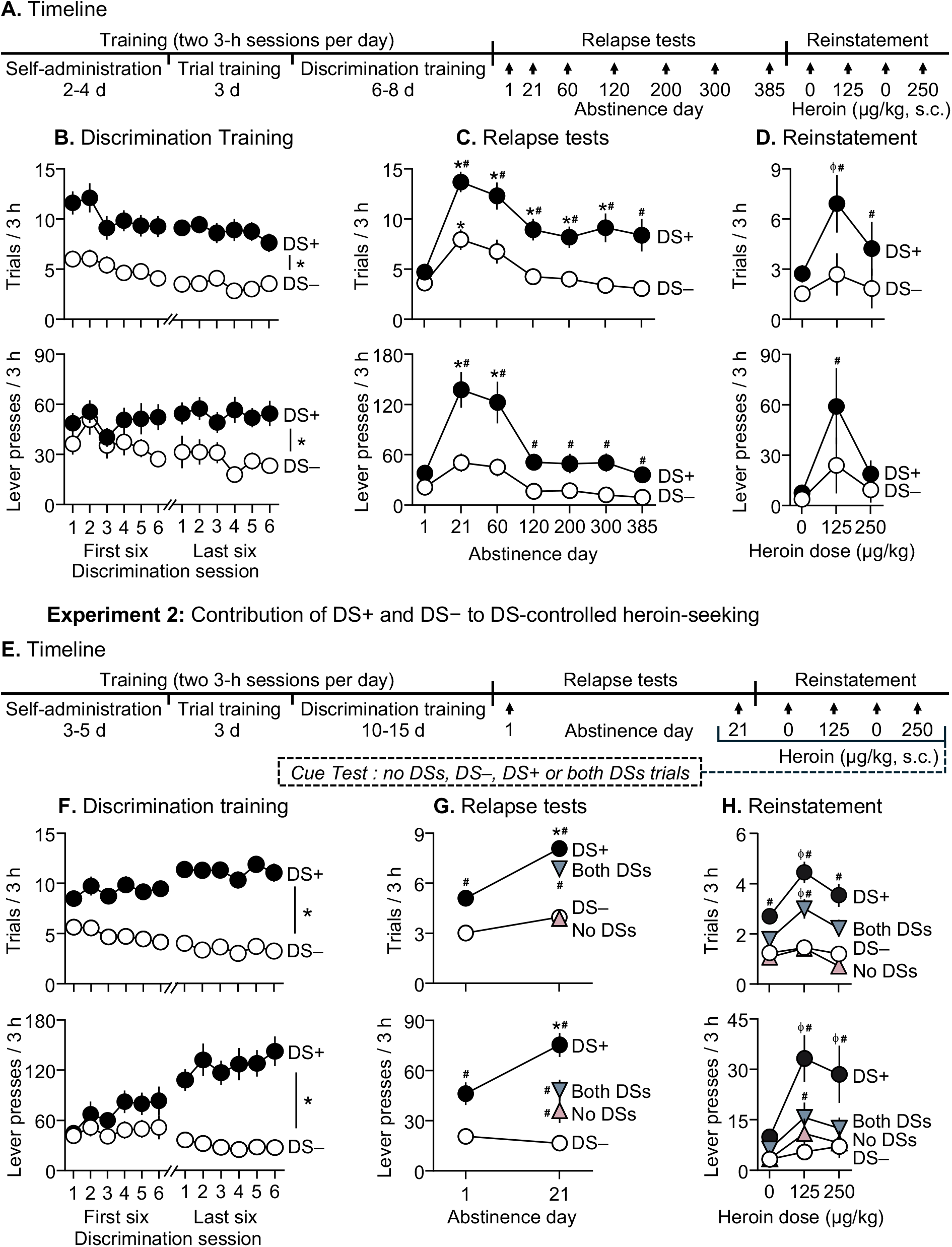
Experiment 1:Incubation of DS-controlled heroin seeking during abstinence. **A**. Experimental timeline for experiment 1 **B**. <u>Discrimination training.</u> Rats first learned to self-administer heroin (0.025 mg/kg/inf.) on a fixed ratio 1 (FR1) reinforcement schedule over 4-8 sessions followed by 2-6 sessions of trial training during which either 30 DS+ or 30 DS− trials were presented during 3-h sessions (data not shown). Next, rats learned to discriminate DS+ from DS− trials within the same session (30 trials each of DS+ and DS− trials presented in pseudorandomized order). Mean (±SEM) number of trials with at least one lever press (top), and lever presses during each trial type (bottom) during a 3-h discrimination training session. * Denotes significant difference (p<0.05) between DS+ and DS− (n=10 M, 10 F). **C**. <u>Relapse tests.</u> Lever presses during DS+ trials peaked on abstinence day 21 and persisted over 385 days. Mean (±SEM) number of trials (top) and lever presses (bottom) during the 3-h relapse test sessions (30 trials each of DS+ and DS− presented in pseudorandomized order) under extinction conditions. * denotes significant (p<0.05) difference from responding during day 1. ^#^ denotes significant (p<0.05) difference between DS+ and DS− responding during the test sessions (day 1 (n=10 M, 10 F), day 21 (n=10 M, 10 F), day 60 (n=10 M, 10 F), day 120 (n=10 M, 10 F), day 200 (n=10 M, 10 F), day 300 (n=7 M, 9 F), day 385 (n=4 M, 9 F). **D**. <u>Reinstatement test.</u> Rats reinstated DS-controlled heroin-seeking in response to injections of heroin (125 and 250 µg/kg, s.c), but not saline. Mean (±SEM) number of trials (top) and lever presses (bottom) during the 3-h saline- or heroin-primed reinstatement test sessions (30 trials each of DS+ and DS− presented in pseudorandomized order) under extinction conditions. ^ϕ^Denotes significant (p<0.05) difference from responding after saline-prime (averaged across the two saline sessions). ^#^ denotes significant difference (p<0.05) between DS+ and DS− responding during the test session (n=4 M, 9 F). **Experiment 2: Individual contributions of DS+ and DS− during DS-controlled heroin seeking** **E**. Experimental timeline for experiment 2 **F**. <u>Discrimination training.</u> Rats first learned to self-administer heroin (0.025 mg/kg/inf.) on an FR1 schedule over 6-10 sessions followed by 6 sessions of trial training during which either 30 DS+ or 30 DS− trials were presented over 3-h sessions (data not shown). Next, rats learned to discriminate DS+ from DS− trials within the same sessions (30 trials each of DS+ and DS− trials in pseudorandomized order). Mean (±SEM) number of trials with at least one lever response (top), and lever-presses during each trial type (bottom) during a 3-h discrimination training session. ^#^ Denotes significant difference (p<0.05) between DS+ and DS− (n=21 M, 15 F). **G**. <u>Relapse tests and DS-control.</u> Rats showed reliable DS-controlled heroin-seeking on abstinence day 1 (data shown for first 15 of 30 trials each of DS+ and DS−, pseudorandomized order, extinction conditions) and showed incubation of DS-controlled heroin-seeking during DS+, but not DS−, on abstinence day 21 (15 each of *no-DSs, DS+, DS−*, and *both-DSs* trials in pseudorandomized order, extinction conditions). Non-reinforced responding during the day 21 test was low during *no-DSs* and *DS−* trials, intermediate during *both-DSs* trials, and highest during *DS+* trials, indicating that DS+ and DS− independently control the expression and suppression of DS-controlled heroin-seeking during abstinence. Mean (±SEM) number of trials with at least one lever response (top), and lever presses during each trial type (bottom) during the first 15 DS+ and DS− trials on day 1, and 15 each of *no-DSs, DS+, DS−*, and *both-DSs* trials on day 21. * Denotes significant (p<0.05) difference from responding during day 1 test (first 15 trials). ^#^ denotes significant difference (p<0.05) from DS− responding during the test session (n=21 M, 15 F). **H**. <u>Reinstatement test and DS-control.</u> Rats reinstated DS-controlled heroin-seeking in response to injections of heroin (125 and 250 µg/kg, s.c), but not saline. Non-reinforced responding was low during *no-DSs* and *DS−* trials, intermediate during *both-DSs* trials, and highest during *DS+* trials, indicating that DS+ and DS− independently control the expression and suppression of DS-controlled drug seeking during reinstatement. Mean (±SEM) number of trials (top) and lever presses (bottom) during the 3-h saline- or heroin-primed reinstatement test sessions (15 trials each of *no-DSs, DS+, DS−*, and *both-DSs* trials presented in a pseudorandomized manner) under extinction conditions. ^ϕ^Denotes significant (p<0.05) difference from responding on the same trial type after saline-prime (averaged across the two saline sessions). ^#^ denotes significant difference (p<0.05) from DS− responding within a test session (n=20 M, 15 F).

### Contribution of DS+ and DS− to DS-controlled heroin-seeking

In Experiment 2 (Figure 1E), we tested whether the two DSs exert opposing control over heroin-seeking during abstinence: DS+ provoking heroin-seeking and DS− suppressing it. After discrimination training as in Experiment 1, we tested rats (n=25 male, 25 female) for DS-controlled heroin-seeking on abstinence day 1, then returned them to their homecages. On day 21, we assessed individual contributions of DS+ and DS− to heroin-seeking. We measured non-reinforced responding during four trial types representing all combinations of DS+ (on/off) and DS− (on/off): no DSs, DS+ only, DS− only, both DSs (15 trials per type). Next, we conducted three additional 3-h extinction sessions and tested heroin-primed reinstatement under the four trial conditions (15 trials per type).

## Results

### Discrimination training (Experiments 1–2)

For each DS trial type in each session, we recorded number of trials with at least one lever-press (“*trials*”) and number of lever presses (“*lever-presses*”) across trials. Rats acquired heroin self-administration (final session mean infusions: 52.8±6.8 and 50.7±6.8 per 3-h in Experiments 1-2) and learned to discriminate the DS+ from the DS− during discrimination training (Fig. 1B, 1F, for full statistical reporting see Supplemental Online Section). We observed no sex differences in heroin self-administration, discrimination performance, or relapse-related measures described below (data not shown).

### Timecourse and persistence of DS-controlled heroin-seeking during abstinence

#### DS-controlled drug-seeking

Rats showed higher heroin-seeking during DS+ trials than during DS− trials across all relapse tests, extending to 385 abstinence days. Heroin-seeking during DS+ trials (and, to a lesser extent, during DS− trials) was greater after 21 and 60 days than after 1 day, indicating incubation of DS-controlled heroin-seeking (Fig. 1C). We analyzed the relapse-test data using Linear Mixed-Effects Model with within-subject factors of Abstinence Day (1, 21, 60, 120, 200, 300, 385) and DS (DS+, DS−). For both measures, we observed significant effects of Day (*trials*: F(2.7,52.1)=14.6, p<0.001; *lever-presses*:F(2.1,40.6)=11.9, p<0.001), DS (*trials*: F(1,19)=34.5, p<0.001; *lever-presses*: F(1,19)=37.9, p<0.001), and Day × DS (*trials*: F(3.2,48.7)=5.0, p=0.0036; *lever-presses*: F(2.9,45.1)=8.3, p<0.001).

#### Heroin priming

Following extinction of DS+ responding, heroin-priming reinstated heroin-seeking during DS+, but not DS−, trials, with a larger effect at the lower dose (Fig. 1D). We analyzed *trials* and *lever-presses* using within-subjects factors of Heroin Dose (Saline [0 mg/kg; mean of saline sessions], 125 µg/kg, 250 µg/kg)and DS (DS+, DS−). For both measures, we observed significant effects of DS (*trials*: F(1,12)=14.6, p=0.002; *lever-presses*: F(1,12)=12.3, p=0.004), and Dose × DS (*trials*: F(1.7,20.9)=6.8, p=0.007; *lever-presses*:F(1.3,15.3)=9.5, p=0.005).

### Contribution of DS+ and DS− to DS-controlled heroin-seeking during abstinence

#### DS-controlled drug-seeking

As in Experiment 1, heroin-seeking during DS+, but not DS−, trials increased between 1 and 21 abstinence days, indicating incubation of DS-controlled heroin-seeking. On day 21, heroin-seeking was low during no-DS and DS− trials, intermediate during both-DS trials, and highest during DS+ trials (Fig. 1G). We first analyzed *trials* and *lever-presses* on abstinence days 1 and 21 (15 trials each of DS+ and DS−) using within-subject factors of Abstinence Day (1, 21) and DS (DS+, DS−). For both measures, we observed significant effects of Day (*trials*: F(1,35)=26.5, p<0.001; *lever-presses*: F(1,35)=8.0, p=0.008), DS (*trials*: F(1,35)=137.5, p<0.001; *lever-presses*: F(1,35)=70.2, p<0.001), and Day × DS (*trials*: F(1,35)=11.3, p=0.002; *lever-presses*: F(1,35)=25.5, p<0.001). To test the independent contributions of DS+ and DS− on day 21, we analyzed *trials* and *lever-presses* for four trial types using within-subject factors of DS+ (on, off) and DS− (on, off). For *trials*, we observed significant effects of DS+ (F(1,35)=114.5, p<0.001) and DS+ × DS− (*trials*: F(1,35)=7.5, p=0.009). For *lever-presses*, we observed significant effects of DS+ (F(1,35)=40.5, p<0.001) and DS− (F(1,35)=12.8, p=0.001).

#### Heroin priming

Following extinction of DS+ responding, heroin-priming selectively reinstated heroin-seeking during DS+ and both-DS trials, but not DS− or no-DS trials (Fig. 1H). We analyzed *trials* and *lever-presses* using within-subject factors of Heroin Dose (0, 125, 250 µg/kg), DS+ (on, off) and DS− (on, off). We observed significant Dose × DS+ (*trials*: F(2,68)=6.8, p=0.002; *lever-presses*: F(2,68)=4.2, p=0.018) and DS+ ×S− (*trials*: F(1,34)=23.3, p<0.001; *lever-presses*: F(1,34)=13.8, p<0.001).

## Discussion

We combined our trial-based DS-controlled drug self-administration model [9] with the classical incubation of drug craving model [3] to examine time-dependent changes and persistent relapse vulnerability across a large part of the adult rat lifespan.

We found that DS-controlled relapse to heroin-seeking persisted for over one year of abstinence, as shown by consistently higher responding during DS+ than DS− trials. Additionally, responding to the DS+ progressively increased (incubated) over the first 3 weeks of abstinence and remained elevated for up to 2 months. These results extend previous rat studies showing persistent responding to discriminative and conditioned stimuli previously paired with psychostimulant (cocaine and methamphetamine) self-administration, where responding increased during early abstinence and persisted for up to 6-10 months [9-12]. While a smaller increase occurred during DS− trials in early abstinence, responding remained significantly lower than during DS+ trials. This modest DS− increase likely reflects response generalization during operant extinction, when the reinforcer is unavailable, rather than a loss of stimulus control [13].

We also found that after over 1 year of abstinence, DS+ and DS− controlled the response to heroin-priming injections: reinstatement occurred during DS+ trials but not DS− trials, consistent with prior findings with cocaine priming [8,14]. Additionally, as with cocaine [15], DS+ and DS− exerted opposing control over heroin-primed relapse: DS+ promoted relapse, whereas DS− inhibited it. Together, these findings underscore the importance of future studies on the neural mechanisms underlying enduring bidirectional DS control over relapse.

The persistence of relapse vulnerability and DS control across the adult rat lifespan has two potential clinical implications. First, the strong and enduring influence of drug-associated cues, such as the DS+ signaling heroin availability used here, suggests that behavioral and pharmacological addiction treatments may need to extend across prolonged abstinence. Second, the sustained inhibitory control of relapse by the DS− signaling drug non-availability for more than a year suggests that behavioral interventions incorporating explicit signals of drug non-availability may hold promise for relapse prevention.

## Author Contributions

RM, YS, and BTH designed the experiments; RM, ORD, DQP, VAL, SJW, JL, AS, ARH, and ON ran the experiments, collected the data, and performed statistical analyses; RM, YS and BTH wrote the paper. All authors reviewed and approved the final version prior to submission.

## Acknowledgements

We thank Dr. Brendan J. Tunstall and Dr. Jennifer M. Bossert for their input during initial stages of the study and members of the Hope and Shaham laboratories for thoughtful comments on the manuscript.

## Data Availability

The complete dataset of the study is available from RM and YS upon request.

