## Supplementary Online Materials for "Persistent vulnerability to heroin relapse across the adult lifespan in rats"

Madangopal et al.

**Table of Contents**

1. Detailed Methods

2. Statistical Table

**Detailed Methods**

**Subjects**

We used male and female Sprague–Dawley rats (Charles River, Raleigh, NC; n = 38 males, 38 females), weighing 200–350 g prior to surgery. Rats were group-housed (two per cage) for at least 1 week before surgery and then individually housed for the remainder of the experiment. We brought rats to the self-administration chambers for training (two 3-h sessions per day) and testing (one 3-h session per day). For all experiments, we maintained the rats under a reverse 12:12 h light/dark cycle (lights off at 8 AM) with free access to standard laboratory chow and water in their home cages. All procedures were approved by the NIDA IRP Animal Care and Use Committee and followed the guidelines outlined in the Guide for the Care and Use of Laboratory Animals [1].

We excluded 25 rats (Experiment 1: n = 3 males, 3 females; Experiment 2: n = 4 males, 10 females) due to loss of catheter patency during training, or failure to acquire discriminated responding. Additionally, not all rats that completed discrimination training underwent the multiple relapse tests due to health-issues during the prolonged abstinence period. The number of rats included in each test is indicated in the corresponding figure legends.

**Drugs**

We received 10 mg/ml diamorphine-HCl (heroin) diluted in sterile saline from the NIDA pharmacy. We chose a unit dose of 0.025 mg/kg per infusion for self-administration training and maintained the same unit dose during discrimination training. We selected this dose to achieve ~30 infusions per 3-h session, comparable to our previous cocaine studies [2,3] in which the unit cocaine dose was 0.75 mg/kg per infusion.

**Surgery**

*Intravenous catheter implantation:* For all behavioral experiments, we implanted rats with silastic catheters using previously described methods [2-4]. Briefly, we anesthetized the rats with isoflurane gas (5% induction, 1-3% maintenance), passed the catheters subcutaneously to the mid-scapular region and then inserted them into the right jugular vein. We attached the catheter tubing to a modified 22-gauge cannula and cemented it in polypropylene mesh implanted under the skin on the rat’s back.

*Post-operative procedures:* We administered ketoprofen (2.5 mg/kg, s.c., Covetrus, formerly Butler Schein) after surgery to relieve pain and allowed rats to recover for 5-7 days prior to drug self-administration training. We flushed the catheters daily with sterile saline containing gentamicin (APP Pharmaceuticals, 4.25 mg/ml) during the recovery and training phases. For rats that lost catheter patency during early stages of training, we inserted a new catheter into the left jugular vein and continued the experiment after 2-3 recovery days. We excluded rats from testing if they lost patency during the last few days of discriminated heroin-taking.

**Behavioral apparatus**

In all behavioral experiments, we trained and tested all rats in Med Associates self-administration chambers (Med Associates ENV-007) enclosed in a ventilated, sound-attenuating cabinet with blacked out windows. The layout of the behavioral chambers was as previously described [2-4]. Each chamber was equipped with a single retractable lever that served as the operand manipulandum, a discriminative stimulus (light, Med Associates ENV-221M) that signaled heroin availability on the left panel and another discriminative stimulus (light, Med Associates ENV-221M) that signaled unavailability of heroin on the right panel of the same side wall, equidistant from the central retractable lever. In addition to location, we used red or white lens caps to differentiate between the two discriminative stimuli and counterbalanced light color across the boxes in both experiments. We used a single speed syringe pump (Med Associates PHM-100, 3.33 RPM) placed outside the sound-attenuating cabinet to deliver intravenous heroin infusions. The pump drove a 20 mL syringe, that was connected (via a liquid swivel) to the rat’s catheter using polyethylene-50 tubing protected by a metal spring.

**Behavioral procedures**

We adapted our published trial-based discriminated cocaine self-administration procedure and trained rats to discriminatively self-administer heroin (0.025 mg/kg/inf.). Training consisting of three phases – heroin self-administration, trial training, and discrimination training. Below we briefly outline these procedures; see [4] for full protocol details.

*Heroin Self-Administration:* We first trained rats to lever press for heroin reward during two 3-h sessions per day separated by 30 min. The start of a session was signaled by the illumination of a light cue on the right side of a central retractable lever, followed 30-s later by the presentation of this lever for 180 min. The light remained on for the duration of the session and served as a discriminative stimulus for heroin reward availability. The same light was later used as a discriminative stimulus to signal availability of heroin during trial-based discrimination training. Throughout the session, responses on this lever were rewarded with an intravenous heroin infusion under a fixed-ratio-1 (FR1) reinforcement schedule. The unit dose of heroin was 0.025 mg/kg/infusion (0.1 ml/infusion) delivered over 3.5 s; drug delivery was not paired with any discrete cues. The infusion duration also served as the timeout period, during which lever presses were recorded but not reinforced. Additional lever presses following this infusion period were also reinforced on the same schedule. At the end of each 3-h session, the discriminative stimulus was turned off and the lever was retracted. We recorded (1) the total number of lever presses and (2) the total number of infusions received during the entire session. We gave rats 4-10 self-administration training sessions to acquire stable responding before switching them to trial training for heroin reward.

*Trial training:* We trained rats in two 3-h trial training sessions per day for 1-3 days. Each session consisted of 30 discrete trials separated by a variable inter-trial interval – the start of each trial was signaled by the illumination of a discriminative stimulus for 30 s, following which rats were given access to the central retractable lever for 60 s. During this initial trial training, each session consisted of only one of two possible trial types – trials in which heroin reward was available (DS+ trials) or trials where heroin reward was not available (DS- trials).

DS+ trials were signaled by the same DS used during continuous access self-administration (light on right side of lever, counterbalanced for red or white light). During DS+ trials, responses on the lever resulted in an intravenous heroin infusion (0.025 mg/kg/infusion delivered over 3.5 s) under an FR1 schedule; drug delivery was not paired with any discrete cues. The infusion duration also served as the timeout period, during which lever presses were recorded but not reinforced. Additional lever presses during this 60-s period (outside the timeout period) were also reinforced on the same schedule. Sixty seconds after lever presentation, the DS+ was turned off and the lever retracted, signaling the end of the trial.

DS- trials were signaled by the other available DS (light on left side of lever, counterbalanced for red or white light). During DS- trials, all responses on the lever were recorded but not reinforced. Sixty seconds after lever presentation, the DS- was turned off and the lever retracted, signaling the end of the trial.

We trained all rats on DS+ trials in the morning session and DS- trials in the afternoon session. We used three behavioral measures to monitor training during this phase – (1) the total number of infusions received during DS+ trials, (2) the total number of DS+ and DS- trials with at least one lever press, and (3) the total number of responses made during DS+ vs. DS- trials during each 3-h session.

*Discrimination training:* We trained rats on the trial-based discrimination procedure for two 3-h sessions per day, separated by 1 h. In each of these sessions, rats received a total of 60 discrete trials; 30 DS+ trials and DS- trials were intermixed and presented in a pseudorandomized order such that rats received no more than two consecutive presentations of the same trial type during the session. For this phase of training, we recorded (1) the total number of infusions received during DS+ trials and (2) the total number of DS+ and DS- trials with at least one lever press, and (3) total number of responses made during DS+ vs. DS- trials during each 3-h session.

*Abstinence phase*: During the abstinence phase for both experiments, we housed rats in individual cages in the animal facility and handled them 1-2 times per week. In Experiment 1, after rats successfully acquired heroin discrimination, we housed them in the vivarium for up to 385 additional days and tested them repeatedly for relapse after progressively longer durations of abstinence from heroin. In Experiment 2, after rats successfully acquired discrimination, we housed them in the vivarium for 20 additional days and examined the individual contributions of DS+ and DS- to persistent DS-controlled heroin seeking on abstinence day 21.

*Relapse tests:* In Experiment 1, after the last discrimination training session, we returned rats to their homecages for a period of experimenter-imposed abstinence and then tested them repeatedly for DS-controlled heroin-seeking on abstinence days 1, 21, 60, 120, 200, 300, and 385. In Experiment 2, we tested rats for DS-controlled heroin-seeking 24 h after the last discrimination training session (i.e., on abstinence day 1) then returned them to their homecages for 20 days of experimenter-imposed abstinence. The experimental conditions during these relapse tests were the same as the trial-based discrimination training session, except that responses on the lever were not reinforced in either DS+ or DS- trials (30 each DS+ and DS- trials, pseudorandomized presentation, extinction conditions). As during discrimination training, we recorded (1) the total number of DS+ and DS- trials with at least one lever press, and (2) total number of responses made during DS+ vs. DS- trials during the 3-h relapse test session. We operationally define relapse as non-reinforced drug seeking following a period of abstinence.

*Four-condition relapse test:* In Experiment 2, we examined the individual contributions of DS+ and DS- to persistent DS-controlled heroin seeking on abstinence day 21 using a modified version of the relapse test described above. We presented all rats with four combinations of DS+ (on, off) and DS- (on, off) in a 2x2 full factorial design - namely *no DSs*, *DS+*, *DS-*, or *both DSs* (15 presentations per type; presented in pseudorandom order) during a 3-h session. Responses on the lever were not reinforced in any of the four trial types. We recorded (1) number of trials with at least one non-reinforced response, and (2) total number of non-reinforced responses made in each trial type during the 3-h relapse test session.

*Heroin-primed reinstatement tests*: In Experiment 1, after the final relapse test (day 385), we gave rats one additional extinction session and then tested the rats for heroin-priming induced reinstatement during four separate sessions, run on consecutive days. We administered a subcutaneous (s.c.) injection of saline or heroin 5 min prior to the start of the test session. We used an ascending dose order for heroin in order (125, 250 µg/kg, within-subjects design) to minimize a carry-over effect of a given priming dose on the subsequent priming dose. We conducted saline-primed reinstatement tests before and between heroin-primed reinstatement tests and averaged the saline-prime data for analysis. The experimental conditions during reinstatement test were the same as during the relapse sessions. We recorded (1) the total number of DS+ vs. DS- trials with at least one lever press and (2) the total number of responses made during DS+ vs. DS- trials over the entire heroin-primed reinstatement test session.

*Four-condition heroin-primed reinstatement tests*: In Experiment 2, after the day 21 relapse test, we gave rats three additional extinction sessions and then tested the rats for heroin-priming-induced reinstatement during four separate sessions. Extinction and reinstatement sessions were run on every other day. As in Experiment 1, we gave the rats an s.c. injection of saline or heroin 5 min prior to the start of the test session, used an ascending dose order, and tested for saline-primed reinstatement before and between heroin-primed reinstatement tests. The experimental conditions during the four-condition reinstatement test were the same as during the four-condition relapse sessions. We recorded (1) the total number of DS+ vs. DS- trials with at least one lever press and (2) the total number of responses made during each trial type over the entire heroin-primed reinstatement test session.

**Experimental design & statistical analyses**

Behavioral data were analyzed using GraphPad Prism (version 11.0.0). We used multifactorial repeated-measures ANOVA or mixed-effects models (to account for attrition and missing values) for statistical analyses, followed by appropriate post-hoc tests as outiled below. When the assumption of sphericity was violated, we applied the Geisser-Greenhouse correction. The alpha level was set at 0.05 (two-tailed). Because some models yielded multiple main effects and interactions, we report only those critical for data interpretation.

*Training:* For both experiments, we first trained rats to self-administer heroin on the trial-based procedure described above. We analyzed the number of “successful” trials (denoted as *trials* and defined as making at least one lever press during a trial) and the total number of lever presses (denoted as *lever-presses* and recorded separately for each DS trial type) during each session. For the analysis of discriminated heroin-taking during training (data not shown), we used repeated measures 2-way ANOVA with within-subject factors of discrimination training Session (last 6 discrimination training sessions) and DS type (DS+, DS-).

*Relapse tests*: In Experiment 1, we analyzed across non-reinforced responding on DS+ and DS- trials using 2-way mixed-effect models with the within-subjects factors of Abstinence Day (1, 21, 60, 120, 200, 300, and 385 days) and DS Type (DS+, DS-), followed by Dunnett’s test for pairwise comparisons between the day 1 relapse test and each of the following days’ relapse tests. We also used Bonferroni test for pairwise comparisons between DS+ and DS- for each relapse test day. In Experiment 2, we used mixed 2-way ANOVA with the within-subjects factors duration of Abstinence Day (1, 21 days) and DS Type (DS+, DS-), followed by Bonferroni test for pairwise comparisons between the day 1 and day 21 for each DS, and between DS+ and DS- on each relapse test day.

*Four-condition relapse test:* In Experiment 2, we analyzed across non-reinforced responding on day 21 across the four types of trials using 2-way ANOVA with within-subject factors of DS+ (on, off), and DS- (on, off), accompanied by Dunnett’s test for pairwise comparisons between DS- and other three trial types during the day 21 test.

*Heroin-primed reinstatement tests*: In Experiment 1, for the tests of priming-induced reinstatement, we analyzed across non-reinforced responding on DS+ and DS- trials using 2-way ANOVA with the within-subjects factors Heroin Dose (0, 125, and 250 µg/kg) and DS Type (DS+, DS-). We used Dunnett’s test for pairwise comparisons between saline and heroin priming doses within each DS trial type. We also used Bonferroni test for pairwise comparisons between DS+ and DS- on each reinstatement test day.

*Four-condition heroin-primed reinstatement tests*: In Experiment 2, for the tests of heroin priming-induced reinstatement, we analyzed across non-reinforced responding across four types of trials using 3-way ANOVA with the within-subject factors of Heroin Dose (0, 125, and 250 µg/kg), DS+ (on, off), and DS- (on, off). We used Dunnett’s test for pairwise comparisons between saline and heroin priming doses within each trial type and for pairwise comparisons between DS- and other trial types for each reinstatement test day.

**References**

1.	National Research Council Committee for the Update of the Guide for the C, Use of Laboratory A. The National Academies Collection: Reports funded by National Institutes of Health. Guide for the Care and Use of Laboratory Animals. Washington (DC): National Academies Press (US) Copyright © 2011, National Academy of Sciences.; 2011.

2.	Madangopal R, Ramsey LA, Weber SJ, Brenner MB, Lennon VA, Drake OR, et al. Inactivation of the infralimbic cortex decreases discriminative stimulus-controlled relapse to cocaine seeking in rats. Neuropsychopharmacology. 2021;46(11):1969–80.

3.	Madangopal R, Tunstall BJ, Komer LE, Weber SJ, Hoots JK, Lennon VA, et al. Discriminative stimuli are sufficient for incubation of cocaine craving. Elife. 2019;8.

4.	Lennon VA, Brenner MB, Weber SJ, Komer LE, Madangopal R. Trial-based Discrimination Procedure for Studying Drug Relapse in Rats. Bio-protocol. 2019;9(23):e3445.

**2. Statistical Table**

| **Figure** | **Experimental phase** | **Behavioral measure** | **Factors in analysis** | **Statistical output** | |
| --- | --- | --- | --- | --- | --- |
| 1B, top right | Discrimination training (Last 6 sessions) | Trials over 3 h (n=10 M, 10 F) | RM-ANOVA (Session x DS) | F-value | P-value |
|  |  |  | Session | F (2.659, 50.53) = 0.6681 | 0.5582 |
|  |  |  | DS | F (1.000, 19.00) = 77.05 | <0.0001 |
|  |  |  | Session x DS | F (3.389, 64.39) = 2.011 | 0.1139 |
| 1B, bottom right | Discrimination training (Last 6 sessions) | Lever presses over 3 h (n=10 M, 10 F) | RM-ANOVA (Session x DS) | F-value | P-value |
|  |  |  | Session | F (2.506, 47.61) = 0.5100 | 0.6448 |
|  |  |  | DS | F (1.000, 19.00) = 20.73 | 0.0002 |
|  |  |  | Session x DS | F (3.026, 57.50) = 1.734 | 0.1697 |
| 1C, panel 1 | Relapse tests | Trials over 3 h (n=4-10 M, 9-10 F) | Mixed-effects model (Day x DS) | F-value | P-value |
|  |  |  | Day | F (2.740, 52.06) = 14.65 | <0.0001 |
|  |  |  | DS | F (1.000, 19.00) = 34.49 | <0.0001 |
|  |  |  | Day x DS | F (3.176, 48.69) = 4.997 | 0.0036 |
| 1C, panel 2 | Relapse tests | Lever presses over 3 h (n=4-10 M, 9-10 F) | Mixed-effects model (Day x DS) | F-value | P-value |
|  |  |  | Day | F (2.137, 40.61) = 11.89 | <0.0001 |
|  |  |  | DS | F (1.000, 19.00) = 37.98 | <0.0001 |
|  |  |  | Day x DS | F (2.941, 45.09) = 8.302 | 0.0002 |
| 1D, panel 1 | Reinstatement tests | Trials over 3 h (n=4 M, 9 F) | RM-ANOVA (Dose x DS) | F-value | P-value |
|  |  |  | Dose | F (1.572, 18.86) = 2.621 | 0.1086 |
|  |  |  | DS | F (1.000, 12.00) = 14.55 | 0.0025 |
|  |  |  | Dose x DS | F (1.744, 20.93) = 6.756 | 0.0071 |
| 1D, panel 2 | Reinstatement tests | Lever presses over 3 h (n=4 M, 9 F) | RM-ANOVA (Dose x DS) | F-value | P-value |
|  |  |  | Dose | F (1.042, 12.51) = 3.632 | 0.0787 |
|  |  |  | DS | F (1.000, 12.00) = 12.27 | 0.0044 |
|  |  |  | Dose x DS | F (1.272, 15.27) = 9.499 | 0.0050 |
| 1F, top right | Discrimination training (Last 6 sessions) | Trials over 3 h (n=21 M, 15 F) | RM-ANOVA (Session x DS) | F-value | P-value |
|  |  |  | Session | F (5, 175) = 1.596 | 0.1636 |
|  |  |  | DS | F (1, 35) = 164.9 | <0.0001 |
|  |  |  | Session x DS | F (5, 175) = 0.5849 | 0.7115 |
| 1F, bottom right | Discrimination training (Last 6 sessions) | Lever presses over 3 h (n=21 M, 15 F) | RM-ANOVA (Session x DS) | F-value | P-value |
|  |  |  | Session | F (5, 175) = 1.441 | 0.2117 |
|  |  |  | DS | F (1, 35) = 52.29 | 0.0001 |
|  |  |  | Session x DS | F (5, 175) = 3.957 | 0.0020 |
| 1G, panel 1 | Relapse tests | Trials over 3 h (n=21 M, 15 F) | RM-ANOVA (Day x DS) | F-value | P-value |
|  |  |  | Day | F (1.000, 35.00) = 26.52 | <0.0001 |
|  |  |  | DS | F (1.000, 35.00) = 137.5 | <0.0001 |
|  |  |  | Day x DS | F (1.000, 35.00) = 11.34 | 0.0019 |
| **Figure** | **Experimental phase** | **Behavioral measure** | **Factors in analysis** | **Statistical output** | |
| 1G, panel 2 | Relapse tests | Lever presses over 3 h (n=21 M, 15 F) | RM-ANOVA (Day x DS) | F-value | P-value |
|  |  |  | Day | F (1.000, 35.00) = 8.031 | 0.0076 |
|  |  |  | DS | F (1.000, 35.00) = 70.23 | <0.0001 |
|  |  |  | Day x DS | F (1.000, 35.00) = 25.49 | <0.0001 |
| 1G, panel 1, day 21 | Day 21 Relapse test (4 trial types) | Trials over 3 h (n=21 M, 15 F) | RM-ANOVA (DS+ x DS−) | F-value | P-value |
|  |  |  | DS+ | F (1.000, 35.00) = 114.5 | <0.0001 |
|  |  |  | DS− | F (1.000, 35.00) = 2.634 | 0.1135 |
|  |  |  | DS+ x DS− | F (1.000, 35.00) = 7.529 | 0.0095 |
| 1G, panel 2, day 21 | Day 21 Relapse test (4 trial types) | Lever presses over 3 h (n=21 M, 15 F) | RM-ANOVA (DS+ x DS−) | F-value | P-value |
|  |  |  | DS+ | F (1.000, 35.00) = 40.53 | <0.0001 |
|  |  |  | DS− | F (1.000, 35.00) = 12.77 | 0.0011 |
|  |  |  | DS+ x DS− | F (1.000, 35.00) = 1.186 | 0.2836 |
| 1H, panel 1 | Reinstatement tests (4 trial types) | Trials over 3 h (n=20 M, 15 F) | RM-ANOVA (Dose x DS+ x DS−) | F-value | P-value |
|  |  |  | Dose | F (2, 68) = 7.717 | 0.0010 |
|  |  |  | DS+ | F (1, 34) = 109.5 | <0.0001 |
|  |  |  | DS− | F (1, 34) = 10.30 | 0.0029 |
|  |  |  | Dose x DS+ | F (2, 68) = 6.785 | 0.0021 |
|  |  |  | Dose x DS− | F (2, 68) = 0.9163 | 0.4049 |
|  |  |  | DS+ x DS− | F (1, 34) = 23.30 | <0.0001 |
|  |  |  | Dose x DS+ x DS− | F (2, 68) = 1.144 | 0.3247 |
| 1H, panel 2 | Reinstatement tests (4 trial types) | Lever presses over 3 h (n=20 M, 15 F) | RM-ANOVA (Dose x DS+ x DS−) | F-value | P-value |
|  |  |  | Dose | F (2, 68) = 5.063 | 0.0089 |
|  |  |  | DS+ | F (1, 34) = 29.64 | <0.0001 |
|  |  |  | DS− | F (1, 34) = 8.465 | 0.0063 |
|  |  |  | Dose x DS+ | F (2, 68) = 4.236 | 0.0185 |
|  |  |  | Dose x DS− | F (2, 68) = 4.057 | 0.0217 |
|  |  |  | DS+ x DS− | F (1, 34) = 13.77 | 0.0007 |
|  |  |  | Dose x DS+ x DS− | F (2, 68) = 1.793 | 0.1743 |
